# Cycling cancer persister cells arise from lineages with distinct transcriptional and metabolic programs

**DOI:** 10.1101/2020.06.05.136358

**Authors:** Yaara Oren, Michael Tsabar, Heidie F. Cabanos, Michael S. Cuoco, Elma Zaganjor, Pratiksha I. Thakore, Marcin Tabaka, Charles P Fulco, Sara A. Hurvitz, Dennis J. Slamon, Galit Lahav, Aaron Hata, Joan S. Brugge, Aviv Regev

**Affiliations:** Klarman Cell Observatory, Broad Institute of MIT and Harvard, Cambridge, MA, USA; Department of Cell Biology, Harvard Medical School, 240 Longwood Ave., Boston, MA, USA; Department of Systems Biology, Harvard Medical School, Boston, MA, USA; Department of Medicine, Massachusetts General Hospital, Boston, MA, USA; David Geffen School of Medicine, University of California, Los Angeles, Jonsson Comprehensive Cancer Center, Los Angeles, CA, USA; Department of Biology, Massachusetts Institute of Technology, Cambridge, MA; Howard Hughes Medical Institute, Chevy Chase, MD

## Abstract

Non-genetic mechanisms have recently emerged as important drivers of therapy failure in cancer (Salgia and Kulkarni, 2018), where some cancer cells can enter a reversible drug-tolerant persister state in response to treatment (Vallette et al., 2019). While most cancer persisters, like their bacterial counterparts, remain arrested in the presence of drug, a rare subset of cancer persisters can re-enter the cell cycle under constitutive drug treatment (Sharma et al., 2010). Little is known about the non-genetic mechanisms that enable cancer persisters to maintain proliferative capacity in the presence of drug. Here, using time-lapse imaging, we found that cycling persisters emerge early in the course of treatment of EGFR-mutant lung cancer cells with the EGFR inhibitor osimertinib. To study this rare, transiently-resistant, proliferative persister population we developed Watermelon, a new high-complexity expressed barcode lentiviral library for simultaneous tracing each cell’s clonal origin, proliferative state, and transcriptional state. Analysis of Watermelon-transduced PC9 cells demonstrated that cycling and non-cycling persisters arise from different pre-existing cell lineages with distinct transcriptional and metabolic programs. The proliferative capacity of persisters is associated with an upregulation of antioxidant gene programs and a metabolic shift to fatty acid oxidation in specific subpopulations of tumor cells. Mitigating oxidative stress or blocking metabolic reprograming significantly alters the fraction of cycling persister cells. In human tumors, programs associated with cycling persisters were induced in malignant cells in response to multiple tyrosine kinase inhibitors. The Watermelon system enabled the identification of rare persister lineages, that are preferentially poised through specific gene programs to proliferate under drug pressure, thus exposing new vulnerabilities that can be targeted to delay or even prevent disease recurrence.

---

To characterize the proliferative dynamics of persisters, we studied the response of PC9 lung cancer cells, which carry an oncogenic mutation in the epidermal growth factor receptor (EGFR), to osimertinib, a third-generation EGFR tyrosine kinase inhibitor. We treated the cells at E_max_ concentration (300μM) (**Fig. S1a**) and used live-cell imaging to quantify the number and timing of division events over the course of treatment, as cells either underwent cell death, arrested, or formed visible colonies (**Fig. 1a**).

**Figure 1.**
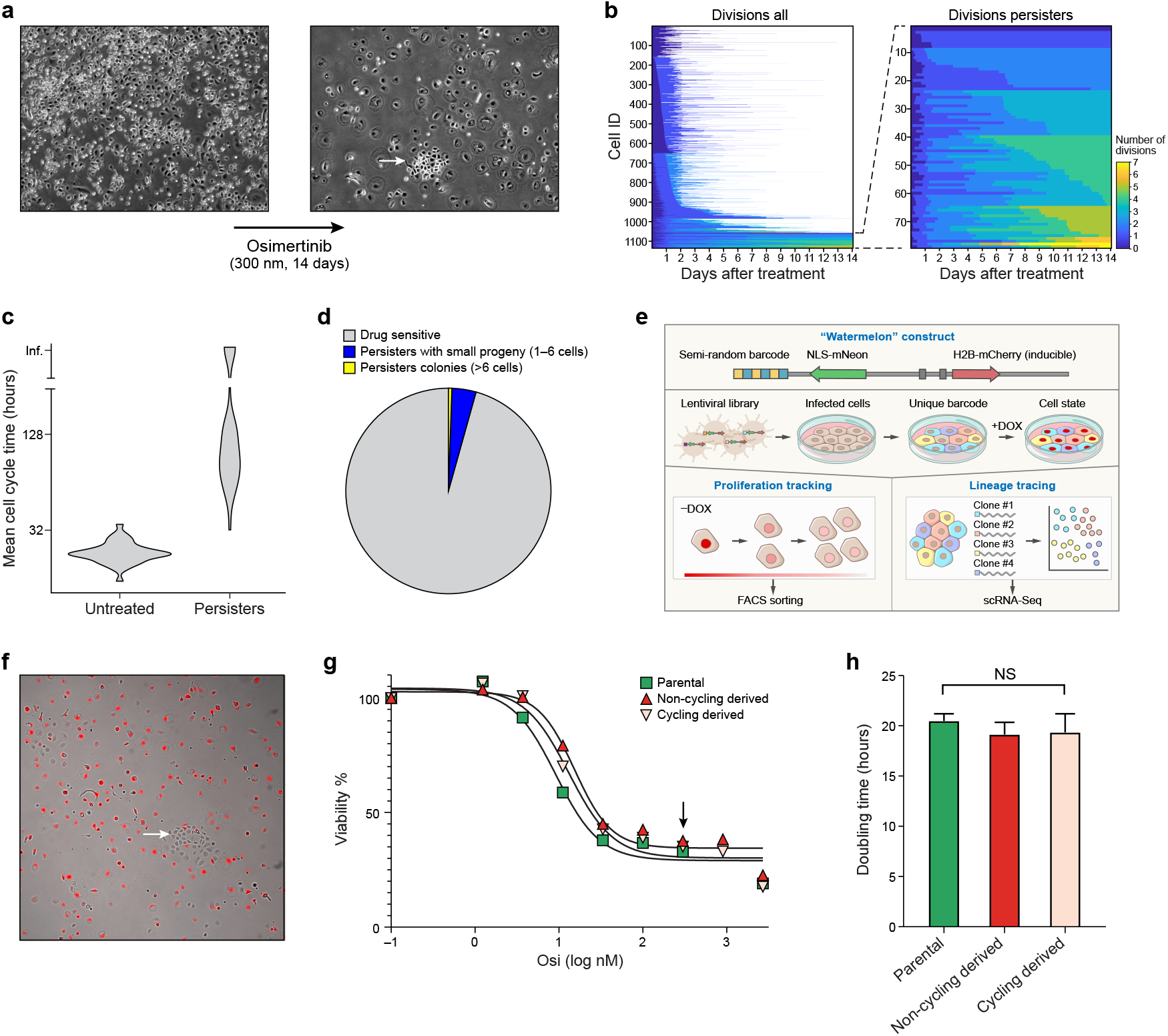
Persister cells contain a rare proliferative subpopulation. ***a***. A colony of cycling persisters. Representative phase contrast images of a colony of PC9 cells before (left) and after (right) 14 days of treatment with 300 nM osimertinib (**Methods**). Arrow: A colony of cycling persisters. Scale bar 100 μm. ***b-d***. Extensive heterogenetiy of cell division profiles of persisters revealed by live cell tracking. ***b***. Left: Number of cell divisions (colorbar) for each of 1,135 individual PC9 cell lineages (rows, left) or only for 77 persister lineages (rows, right) tracked by live imaging along a 14 days osimertinib treatment time course (*x* axis). White: lineage perished. ***c***. Distribution of mean cell cycle time (*y* axis) of untreated and persister cell lineages (*x* axis) from live cell imaging tracking data. ***d***. Proportion of lineages with a large (>6 cells; yellow) or small (1-6 cells, blue) number of progeny or with no progeny (drug sensitive, grey) by the end of the 14 days osimertinib treatment. ***e,f***. The Watermelon system for simultaneous tracing of clonal lineages and transcriptional and proliferative state of individual cells. ***e***. Watermelon vector. Semi-random barcode linked to the NLS-mNeon gene allows for lineage tracing. The doxycycline inducible H2B-mCherry facilitates proliferation tracking via fluorescent dilution. ***f***. Watermelon-PC9 persister cells following 14 days of osimertinib treatment during which dox was included for the first 3 days. Arrowhead: colony of cycling persisters. Fluorescence (red) and phase contrast channel merge is shown. Scale bar 100 μm. ***g***. Both cycling and non-cycling persister cells rapidly reacquire drug sensitivity. Percent of viable cells (*y* axis) from populations derived from parental (green), cycling persister (pink) and non-cycling persister (red) cells treated with different osimertinib concentrations (*x* axis) for 72hr (**Methods**). Black arrow: 300nM, the osimertinib concentration used for establishing persister cells. ***h***. Similar population doubling times for cells derived from persister-cycling and persister-non-cycling subpopulations. Mean doubling time (*y* axis) in populations derived from parental (green), cycling persister (pink) and non-cycling persister (red) cells, grown in drug free media. Error bars: SD. n= three biologically independent experiments. NS, not significant (P > 0.05); two-tailed t-tests.

Only 8% of cell lineages gave rise to persisters, defined as cells that were alive at day 14 of drug treatment (**Fig. 1b**). Cell division profiles of persisters revealed extensive heterogeneity, from persisters that did not divide at all to those that underwent seven divisions (**Fig. 1b**). This was in contrast to the unimodal cell cycle time distribution of the untreated parental cell population (**Fig. 1c**). In line with previous reports that colony-forming persisters are rare (Hata et al., 2016; Sharma et al., 2010), less than 0.5% of the initial cell population (13% of the persister population) gave rise to multi-cellular persister colonies of more than six cells (**Fig. 1d**). Thus, persister cells contain a rare proliferative subpopulation that emerges early in the course of drug treatment.

Identifying mechanisms that allow cells to survive drug treatment and regain proliferative capacity requires measuring the cellular and molecular properties of these rare cells prior to and during drug treatment. To this end, we developed Watermelon, a high complexity lentiviral barcode library, with both green and red fluorescent reporters, for simultaneous tracing of clonal lineages as well as the transcriptional and proliferative state of each cell in the population. Lineage tracing was achieved by mapping a clone-specific expressed DNA barcode in the 3′ untranslated region of an mNeonGreen protein, and proliferation history was monitored by the dilution of a doxycycline (dox)-inducible H2B-mCherry transgene (**Fig. 1e** and **Fig. S1b-e**). We generated a Watermelon library of more than five million barcodes and used it to transduce 10,000 PC9 cells with a Multiplicity of Infection (MOI) of less than 0.3 to minimize barcode overlap between clones (**Methods**).

We tested whether proliferative persisters arise due to a stable, intrinsically lower sensitivity to EGFR inhibition, by comparing the osimertinib sensitivity of cycling and non-cycling persister-derived populations following a short re-sensitization period. We treated Watermelon-PC9 cells with osimertinib for 14 days, with doxycycline (dox) added to the media for the first three days, followed by 11 days of dox chase, during which mCherry was diluted only in proliferating cells, sorted cycling and non-cycling persister cells of day 14 (**Fig. 1f**), and propagated each subset in drug-free media for 20 additional passages. Both cycling and non-cycling persister populations reacquired drug sensitivity rapidly, suggesting that a reversible, rather than a heritable, mutational mechanism underlies the ability to cycle under continuous drug treatment (**Fig. 1g**). Furthermore, cells derived from persister-cycling and persister-non-cycling subpopulations had similar population doubling times, suggesting that the drug-observed phenotypic differences are not due to pre-existing proliferative heterogeneity (**Fig. 1h**).

To identify mechanisms underlying the ability of persister cells to proliferate, we profiled the expression of 56,419 individual Watermelon-PC9 persister cells by scRNA-seq, at four time points (days 0, 3, 7 and 14) along 14 days of osimertinib treatment (**Fig. 2a** and **Fig. S2a-c**). In particular, on day 14 we sorted persister cells by mCherry expression into three sets: cycling (mCherry^low^), moderately cycling (mCherry^medium^) and non-cycling (mCherry^high^) persisters, and profiled each of these by scRNA-seq separately. We used published signatures(Tirosh et al., 2016) to assign each cell to a specific cell phase (**Fig. 2b)**, ascribed cells to lineages based on the expressed barcodes detected by scRNA-seq (**Fig. 2c** and **Fig. S2d**), and related cells by their profiles using a force-directed layout embedding (FLE) (**Fig. 2d**).

**Figure 2.**
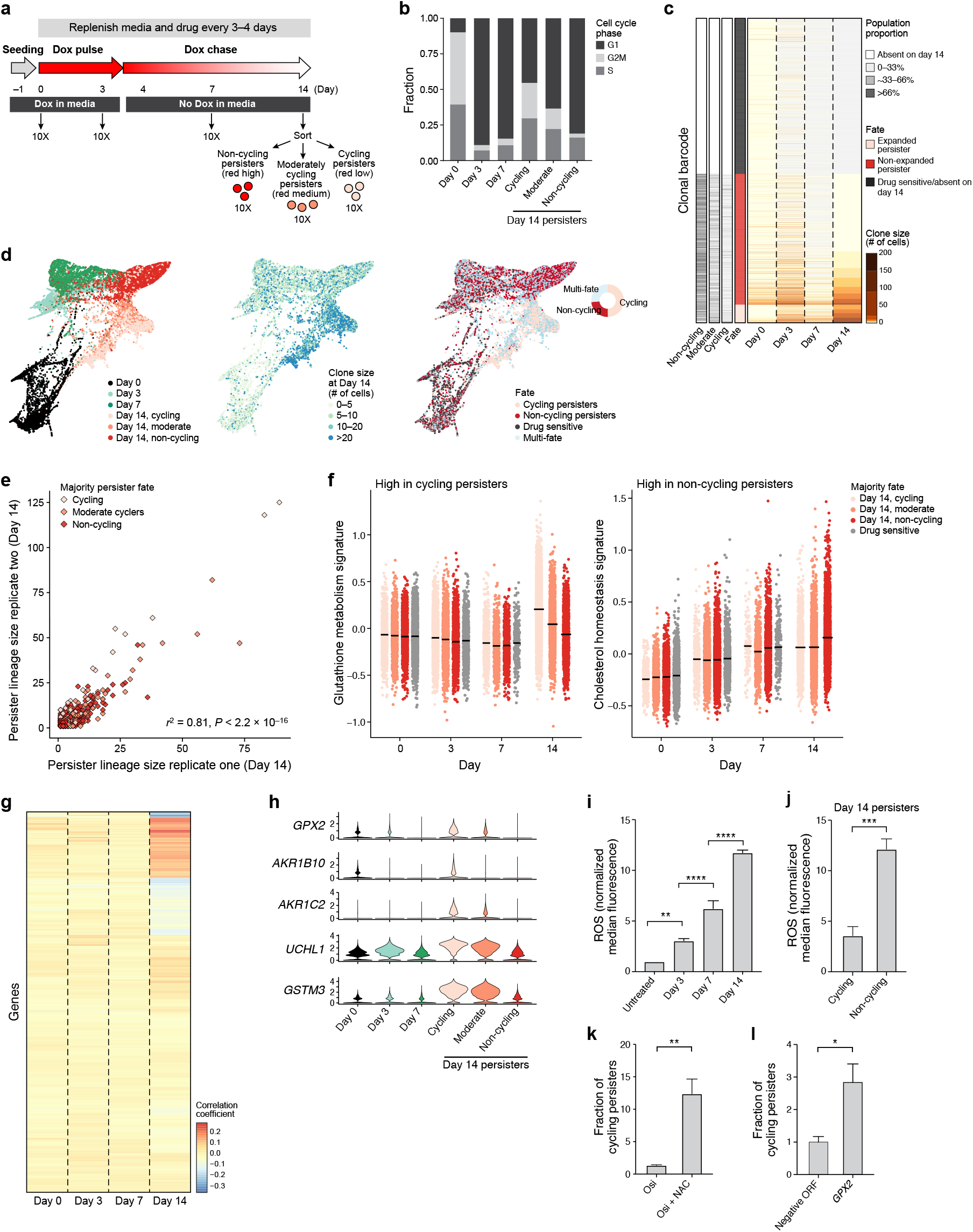
Cycling and non-cycling persisters arise from different cell lineages that express distinct transcriptional programs. ***a***. Experimental scheme. Vertical arrows: scRNA-seq collection time point. ***b***. Distinguishing persisters by proliferative history. Proportion of cells (*y* axis) in each cell cycle phase (colored stacked bars) based on cell cycle scores inferred from scRNA-seq data across time points (*x* axis). ***c***. Ascribing cells to lineages and their fate along the time course. Lineage size (red/yellow colorbar) for each lineage barcode (rows) across time points. Lineages are sorted by fate at day 14 (red/black/pink bar), and marked by the proportion of each of the mCherry populations on day 14 in that lineage (three left most bars). ***d***. Cycling and non-cycling cells follow distinct trajectories. Force directed layout embedding of scRNA-seq profiles (dots) colored by time point and day14 mCherry expression (left); day 14 clone size (middle) or fate (right) of the lineage the cells belong to. Inset, right: fate distribution at day 14. ***e***. Persister clones have reproducible proliferative potential under drug. Clone size on day 14 of each persister lineage barcode inferred from scRNA-seq (*x, y* axes) in two independent experiments seeded from the same barcoded founding cell population. Color: Persister fate of majority of cells in the lineage of combined replicates. Bottom right: Pearson correlation coefficient. ***f-h***. Distinct transcriptional programs in cycling and non-cycling persisters. ***f***. Signature scores (y axis, **Methods**) of glutathione metabolism (left) and the cholesterol homeostasis signaling (right) pathways in cells profiled at each time point (*x* axis) stratified by their lineage majority fate at day 14 (color legend). ***g***. Pearson correlation coefficient (colorbar) of each gene’s (row) expression in each persister cell in a given timepoint (columns) and the cells’ corresponding lineage size in day 14. ***h***. Distribution of expression levels (log_2_(TPM+1), *y* axis) at each time point and population (*x* axis) for 5 of the top 10 genes positively correlated with persister day 14 lineage size. ***i-l***. Role for redox balance in regulating the proliferative capacity of persisters. ***i,j***. Mean ROS levels (*y* axis), measured by FACS of CellROX in PC9 cells treated with 300 nM osimertinib across time points (i) or in persister subpopulations on day 14 (j) (*x* axis). ** − P< P < 0.01; **** − P < 0.000; two-tailed t-tests. Error bars: SD. ***k,l***. Mean fraction of cycling persisters (*y* axis) following 14 days of 300 μM osimertinib treatment in (k) cells treated with or without 5mM NAC during days 3-14 (*x* axis), or (l) in PC9 cells stably transfected with control vector or a vector expressing GPX2 protein (*x* axis). Error bars: SD, * P < 0.05; ** P < 0.01; two-tailed t-tests. n= three biologically independent experiments (i− l).

In line with previous reports(Sharma et al., 2010; Taniguchi et al., 2019), osimertinib induced a G1 arrest in day 3 and 7 of treatment. At day 14, cycling persisters (as defined at the end point by mCherry dilution) had higher expression-based cell cycle scores compared with moderate cyclers and non-cycling cells (55%, 36%, 18% cells in G2/M or S phase for cycling, moderate cyclers and non-cycling persisters respectively, **Fig. 2b** and **Fig. S2e**), validating Watermelon’s ability to distinguish persisters by proliferative history. Consistent with our imaging-based analysis, less than 12% of barcodes that were present in persisters at day 14 were part of a large clonal expansion, increasing in size by 5-fold or more during the course of the 14-day assay (**Fig. 2c** and **Fig. S2e**, “expanded persisters”).

The scRNA-seq profiles indicated a gradual change of cell state following drug treatment, with cycling and non-cycling persister cells following distinct transcriptional trajectories, and a subset of cycling persisters resembling untreated cells (**Fig. 2d** left; overlapping light pink and black dots). Examination of clone size dynamics during the course of treatment revealed a time-dependent clonal expansion with the largest increase in clone size observed in the cycling persisters (175, 110 and 53 maximum cells per lineage for cycling, moderate cyclers and non-cycling persisters, respectively, **Fig. 2d**, middle, **Fig. S2f**). Importantly, almost two-thirds of clones were uni-fate, giving rise to either only cycling or only non-cycling persisters, and multi-fate lineages were far less frequent than expected by chance (P<0.001, permutation test, **Fig. 2d**, right).

We hypothesized that this restricted cell fate pattern arises because cells are often committed to a given proliferative fate prior to drug treatment. To test this hypothesis, we compared persister lineage size in two independent replicate experiments using the same starting Watermelon-transduced PC9 cell population, treated with osimertinib for 14 days, sorted to three subpopulations by mCherry expression and profiled by scRNA-Seq. The sizes of individual clones at day 14 were highly correlated between replicates (Pearson *r*^2^ =0.81 P value <2.2*10^−16^, **Fig. 2e**), suggesting that each persister clone has a distinct proliferative potential under drug.

To identify cellular programs that are associated with the ability of persisters to cycle, we searched for gene signatures that are differentially expressed between the persister subpopulations, but are cell cycle independent. In line with previous reports (Hangauer et al., 2017; Raoof et al., 2019), epithelial-mesenchymal transition (EMT) genes and GPX4, a hydroperoxidase that protects cells against membrane lipid peroxidation, were induced by the EGFR inhibitor; however, the levels of induction were similar in both the cycling and non-cycling persister populations (**Fig. S3a, b**), suggesting that these programs are not underlying the ability of persisters to cycle. Cycling persisters exhibited higher expression of glutathione metabolism and NRF2 signatures than non-cycling persisters, whereas cholesterol homeostasis, interferon alpha and Notch signaling signatures were higher in the non-cycling persister subpopulation (**Fig. 2f**, **Fig. S3c, d** and **Supplementary Table 1**). The signatures associated with cycling persister fate were not upregulated prior to drug treatment, suggesting that cycling persisters arise from cells poised to induce these programs, rather than from a selection of cells that already express them prior to drug treatment.

To explore further the transcriptional differences at the lineage level, we correlated gene expression over time with persister clone size at day 14. While at early time points no genes were strongly correlated with persister clone size, expression of a subset of genes at day 14 strongly correlated with persister clone size (**Fig. 2g)**. Of the top ten correlated genes more highly expressed in cells from larger clones, four (AKR1B10, AKR1C2, GPX2, and UCHL1) are targets of the oxidative stress-induced transcription factor NRF2 (**Fig. 2h**), further supporting a role for antioxidant defense programs in persister proliferative fate.

Given the strong relationship between antioxidant transcriptional signatures and persister proliferative capacity, we tested whether prolonged osimertinib treatment induces reactive oxygen species (ROS) and if alleviating ROS enhances cycling persisters. Analysis of osimertinib-treated cells using CellROX, a fluorescent ROS reporter, revealed a strong time-dependent increase in ROS levels (**Fig. 2i**). In line with the observed transcriptional differences between the two persister subpopulations, day 14 cycling persisters exhibit less than a third of the ROS levels measured in the non-cycling population (**Fig. 2j**). Moreover, when we alleviated ROS by treatment with NAC, a ROS scavenger, from day 3 onwards **(Fig. 2k)**, or by stable overexpression of the GPX2 open reading frame (**Fig. 2l**), the fraction of cycling persisters increased significantly. This supports a role for redox balance in regulating the proliferative capacity of persisters.

Because redox balance is tightly linked to metabolism(Reczek and Chandel, 2017), we performed LC-MC/MS metabolic profiling to examine differences between persister subpopulations. Principal component analysis (PCA) over the 229 quantified metabolites separated the cycling persisters, non-cycling persisters and untreated populations (**Fig. 3a** and **Fig. S4a, b**), with 56 metabolites differentially abundant between samples (**Fig. 3b**). In particular, carnitine-linked fatty acids, which are substrates of mitochondrial β-oxidation, were significantly more abundant in the cycling persisters than in the non-cycling persisters (**Fig. 3c**). We examined the oxidation of radiolabeled palmitic acid to ^3^H_2_O to assess fatty acid oxidation (FAO), and indeed found a time-dependent increase in FAO with osimertinib treatment (**Fig. 3d** and **Fig. S5a, b**). Thus, the higher abundance of acylcarnitine species in cycling persisters and the increase in FAO over time with osimertinib treatment both suggest that mitochondrial FAO may contribute to the cycling persister phenotype.

**Figure 3.**
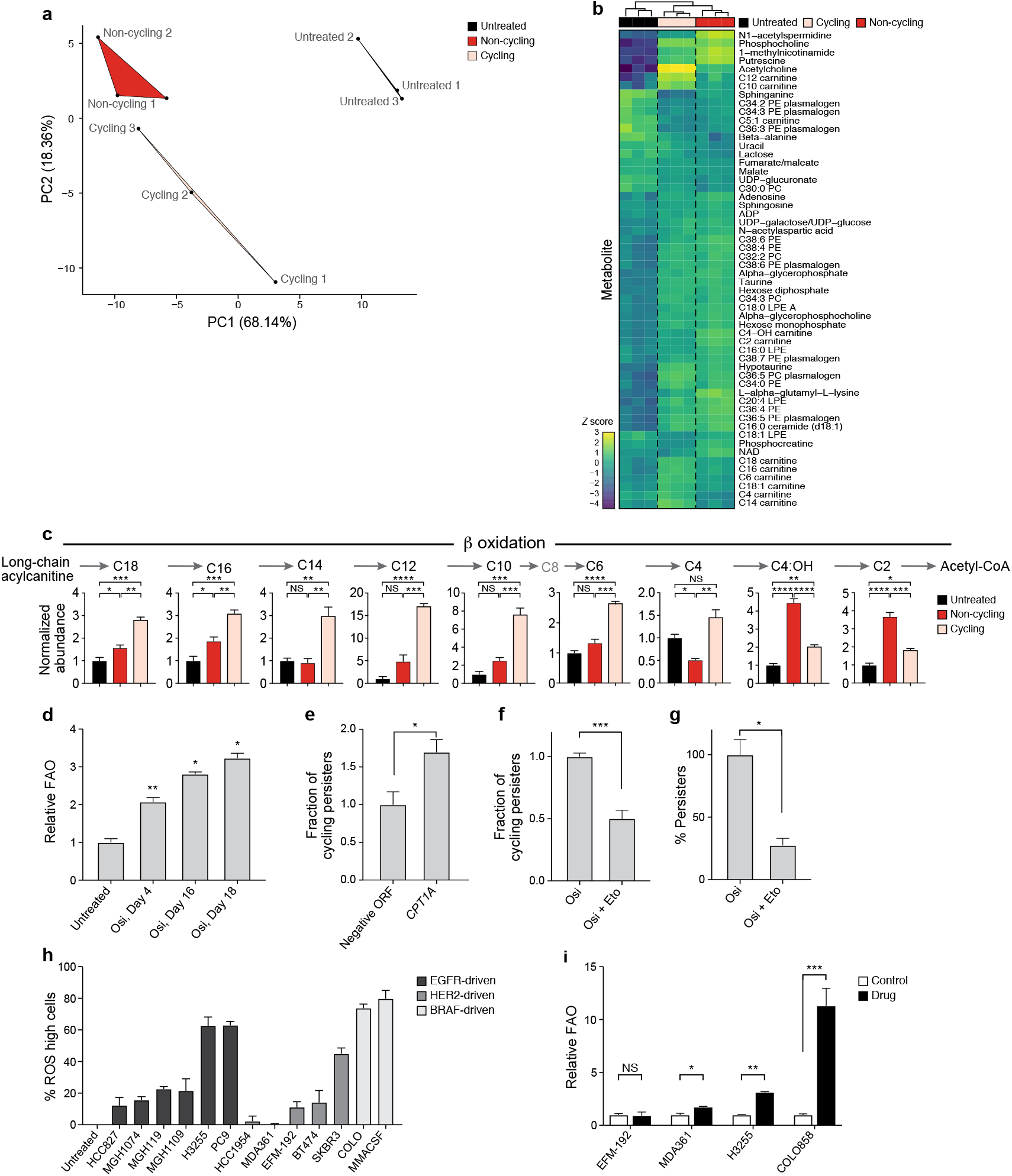
Persister cells shift their metabolism to fatty acid oxidation. ***a,b***. Distinct metabolic profiles of cycling persisters, non-cycling persisters and untreated parental cells. ***a***. The first two principle components (PCs, x and y axis) of a Principal Component Analysis (PCA) of LC-MS/MS metabolite profiles for cycling persisters (pink), non-cycling persisters (red) and untreated parental cells (black). ***b***. Z-scores (colorbar) of metabolites (rows) that are differentially abundant (ANOVA P-value<0.05) between cycling persisters (pink), non-cycling persisters (red) and untreated parental cells (black) (columns). ***c***. β-oxidation metabolites are more abundant in cycling than non-cycling persisters. Mean abundance (*y* axis, normalized relative to untreated samples) of β-oxidation metabolites measured by LC-MS/MS in cycling persisters (pink), non-cycling persisters (red) and untreated parental cells (black) [n=3]. ***d***. Time-dependent increase in FAO with osimertinib treatment. Mean fatty acid oxidation (FAO) levels (y axis, relative to mean of untreated cells) measured by 3H-palmitic acid oxidation in PC9 osimertinib-treated cells over time (*x* axis). ***e-g***. Modulating the FAO pathway affects the proliferative capacity of persisters. ***e***. Mean fraction of cycling persisters (*y* axis) at day 14 of 300 nM osimertinib treatment of PC9 cells transfected with control vector or a vector expressing CPT1A protein (*x* axis). ***f,g***. Mean fraction of cycling persisters (*y* axis, f) and overall fraction of persister cells (*y* axis, g) at day 14 of 300 nM osimertinib treatment alone or with 100μM Etomoxir at days 3-14 (*x* axis). ***h,i***. ROS and FAO increased by drug treatment in persisters of diverse cancer types. ***h***. Mean fraction cells (y axis, %) with higher ROS levels than the corresponding untreated cell line, in lung (black), breast (dark grey) and melanoma (light grey) cell lines treated with 300 nM osimertinib (14 days), 1 μM lapatinib or 1 μM dabrafenib (10 days), respectively. ***i***. Mean FAO levels (*y* axis) relative to untreated controls (white bars) in breast (EFM-192 and MDA361), lung (H3255) and melanoma (COLO855) cell lines treated with 1μM lapatinib, 300nM osimertinib or 1mM dabrafenib, respectively, for 10 days (breast and melanoma) or 14 days (lung). Error bars: SD, from three (d– i) biologically independent samples. * P < 0.05; * * P < 0.01; * * * P < 0.001; * * * *P < 0.0001; two-tailed t-tests.

We next tested if modulating the FAO pathway affects the proliferative capacity of persisters. Overexpressing CPT1A, a rate limiting enzyme that facilitates the transfer of fatty acids into the mitochondria, in Watermelon-PC9 cells resulted in >50% increase in the fraction of cycling persisters following 14 days of osimertinib treatment (**Fig. 3e**). Conversely, blocking FAO with etomoxir, a CPT1 inhibitor (albeit with additional targets), at day 3 of osimertinib treatment, when most sensitive cells have already died, reduced the proliferative capacity of persister cells. Co-treatment with etomoxir for 11 days, at a concentration that has minimal inhibitory effects on untreated cells (**Fig. S5c**), significantly reduced the fraction of cycling persisters (**Fig. 3f**), and the overall fraction of persister cells (**Fig. 3g**). Taken together, these results support our hypothesis that FAO plays a role in establishing the proliferative persister state.

To investigate whether these findings are relevant to other persister models, we chronically treated multiple EGFR-driven lung, HER2-driven breast cancer and BRAF-driven melanoma cell lines with osimertinib, lapatinib and dabrafenib, respectively, and assessed ROS and FAO induction. Three of five breast cancer cell lines and all lung and melanoma cell lines examined showed an increase in ROS following prolonged drug treatment (**Fig. 3h**). In three of the four cell lines, in which a sufficient number of persister cells could be recovered for metabolic labeling, FAO was significantly increased compared with the untreated control (**Fig. 3i**). This suggests that the persister state is supported by a metabolic switch in response to multiple tyrosine kinase inhibitors in cancer cells from different solid tumor types.

Finally, we explored the relevance of these metabolic states in patient tumors, by analyzing single cell RNA-seq profiles from human EGFR-driven lung adenocarcinomas tumor samples from a recent study(Ashley Maynard, 2019) (**Supplementary Table 2**), spanning treatment naïve tumors (TN), residual disease during targeted therapy response (residual disease, RD), and upon establishment of drug resistance (progression, PD). In line with our *in vitro* findings, both fatty acid metabolism (FAM) and ROS pathway signatures were significantly and gradually increased upon treatment from TN to RD to PD (**Fig. 4a, b** and **Fig. S6a, b**).

**Figure 4.**
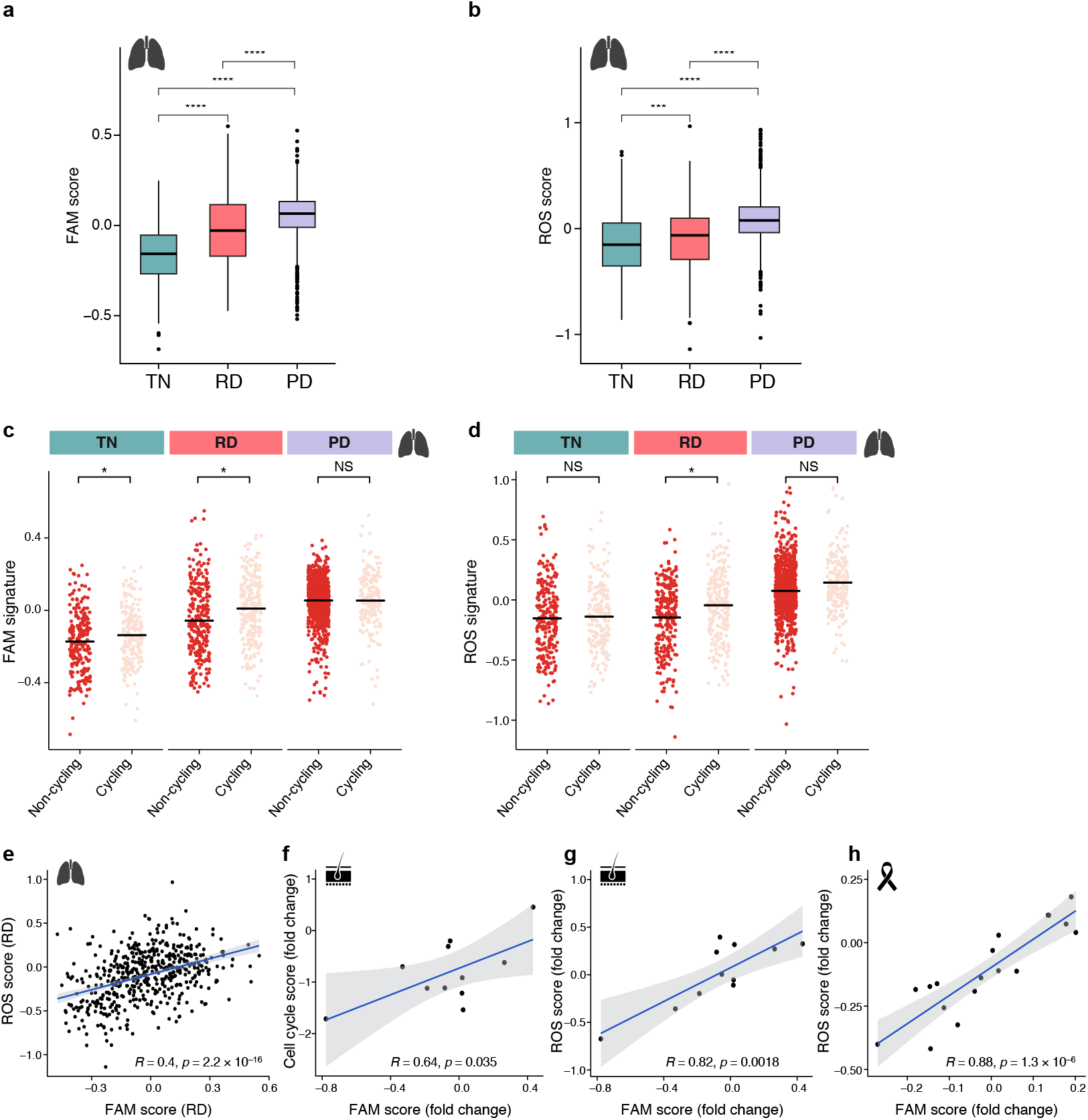
Metabolic shift in tumors treated by oncogene-targeted therapy. ***a,b***. Fatty acid metabolism and ROS pathway signatures increase with treatment in human lung adenocarcinoma tumors. Signature scores for fatty acid metabolism (FAM, a) and reactive oxygen species (ROS, b) in treatment naïve (TN), residual disease (RD) and progressive disease (PD) of human lung adenocarcinoma. *** P < 0.001**** P<0.001; Wilcoxon test. Expression data from NCT03433469 trial. ***c,d***. Higher increase in FAM and ROS signatures in cycling *vs*. non-cycling persister cells. FAM (c) and ROS (d) signature scores (y axis) in cells (dots) stratified by cycling (pink) *vs*. non-cycling (red) status across treatment phases (*x* axis) of lung adenocarcinoma samples. * P < 0.05 (mixed linear model; **Methods**). ***e***. ROS and FAM signatures are correlated in residual disease samples. ROS and FAM signature scores (x and y axes) in cells (dots) from residual disease. Bottom right: Pearson correlation coefficient. ***f-h***. Correlation between ROS and FAM scores in melanoma and breast cancer residual disease. ***f***. Cell cycle (y axis) and FAM (x axis) signature induction score in BRAFi/MEKi-treated melanoma (NCT01072175 trial). ***g,h***. ROS (*y* axis) and FAM (x axis) signature scores in in BRAFi/MEKi-treated melanoma (g) and HER2^+^ breast cancer patients treated with lapatinib (h, NCT00769470 trial). Bottom right: Pearson correlation coefficient.

The increase in ROS and fatty acid metabolism gene expression is associated with proliferation, and is higher in cycling *vs*. non-cycling persister cells (**Fig. 4c, d**). Specifically, when we stratify cells as cycling *vs*. non-cycling by the expression of a set of known cell cycle genes(Patel et al., 2014), cycling persister cells in the RD phase had higher expression of FAM and ROS signatures compared with non-cycling RD cells (P value <0.05 for both pathways after controlling for patient and cell complexity, see Methods, **Fig. 4c, d**). Importantly, ROS and FAM signatures were more strongly correlated in treated than in TN samples (R=0.4, P value=2.2*10^−16^ for RD *vs*. R=0.1, P value=0.01 for TN, **Fig. 4e** and **Fig. S6c**), supporting the importance of oncogene inhibition in linking these pathways.

To test whether similar states are observed in response to other tyrosine kinase inhibitors, we analyzed bulk RNA-seq profiles from two additional tumor types: BRAF-driven melanoma collected at baseline or after up to 22 days of BRAF inhibitor + MEK inhibitor treatment(Kwong et al., 2015), and HER2-driven breast cancer from patients at baseline or after treatment with lapatinib for 2-3 weeks (**Supplementary Table 2)**. In melanoma, cell cycle signature scores were correlated with FAM in treated samples (R=0.64, P value=0.03, **Fig. 4f**) and FAM and ROS induction was strongly correlated only upon treatment (Pearson *r*= 0.8, P value=0.002, **Fig. 4g**, **Fig. S6d**), with 8/11 patients exhibiting induction of at least one of the signatures following treatment. In HER2+ breast cancer, FAM and ROS signature induction were strongly correlated only upon drug treatment (R=0.88 P value=1.3e-6, **Fig. 4h** and **Fig. S6e**), with 50% of patients exhibiting induction of at least one of the signatures following treatment. This supports our model that cycling persisters undergo a metabolic shift in patient tumors in response to tyrosine kinase inhibitor (TKI) therapy.

The understanding that non-mutational mechanisms may play a role in tumor relapse has prompted multiple studies focused on identifying factors that contribute to overall persister fitness(Shaffer et al., 2017; Shah et al., 2019; Viswanathan et al., 2017). However, since most persisters remain arrested during drug treatment (Sharma et al., 2010), factors that contribute to persister-driven relapse are hard to discern by bulk profiling. Watermelon’s ability to simultaneously map the lineage, proliferative history and transcriptional state of individual cells allowed us to identify metabolic and expression adaptations that may facilitate cell cycle re-entry in a rare subset of persister cells. In particular, our finding that some lineages are more poised to undergo a proliferation-promoting adaptive response is in line with studies showing that a subset of cancer cell genomes have a chromatin configuration that renders them more likely to transition to an oncogene-independent state following treatment(Guler et al., 2017; Liau et al., 2017). Taken together, the results presented here illustrate an approach to understand not only what underlies the ability of persisters to cycle but also what drives other clinically important persister traits such as time-to-relapse, thus providing an important step towards the development of therapies that delay disease recurrence.

## Acknowledgments

We thank the Broad Cytometry Facility (P. Rogers, S. Saldi, C. Otis and N. Pirete); Leslie Gaffney and Ania Hupalowska for help with figure preparation and artwork; and Angie Martinez Gakidis for scientific editing.

## Author contributions

Study design, data interpretation and preparation of the manuscript: YO, JSB and AR. Execution of experiments: YO, MT, HFC, MSC, EZ, PIT. Computational analysis: YO, MSC, and MT. Resources: MT, CPF, SAH, DJS, GL, AH.

## Data and materials availability

Raw gene expression data and processed counts are available on GEO, accession no. GSE150949. Code will be available upon request.

