## Supplementary material for "Cycling cancer persister cells arise from lineages with distinct transcriptional and metabolic programs": Material and Methods and supplemental figures

##### Cell culture

EGFR-mutant non-small-cell lung cancer cells PC9 and H3255 (Hata lab) and HER2-amplified breast cancer cell lines MDA361, HCC1954, EMF192 and SKRB3 were cultured in phenol red free RPMI-1640 medium (ThermoFisher Scientific) supplemented with 10% fetal bovine serum (FBS). BT474 breast cancer cell line was grown in phenol red free DMEM/F12 (ThermoFisher Scientific) supplemented with 10% FBS. All breast cancer cell lines were obtained from the Brugge lab. Melanoma cell lines used in this study were obtained from the the Harvard Medical School (HMS) Laboratory of Systems Pharmacology (LSP). COLO858 cell line was grown in phenol red free RMPI 1640 supplemented with 5% FBS and 1% sodium pyruvate (Gibco). MMACSF was grown in DMEM/F12 supplemented with 5% FBS and 1% sodium pyruvate. All cell lines were with supplemented with penicillin and streptomycin (Thermo Scientific). Each cell line was maintained in a 5% CO2 atmosphere at 37 °C. Cell line identities were confirmed by STR fingerprinting and all were found to be negative for mycoplasma using the Universal Mycoplasma Detection Kit (ATCC).

##### Persister cell derivation and treatments

Persister cells were derived using an IC90 drug concentration from treatment of EGFR-mutant non-small-cell lung cancer, BRAF mutated melanoma or HER2-amplified breast cancers with 300uM osimertinib, 1 µM dabrafenib or 1 µM lapatinib respectively, for 10 days for the breast and melanoma cells and 14 days for the lung cells. Fresh media with drug was added every 3-4 days. For watermelon vector induction, doxycycline was added to the media two days prior to drug treatment and was maintained in the media for the first 3 days of drug treatment. Unless otherwise noted, any additional drugs were added to three days treated cells that were maintained under constant drug exposure throughout subsequent treatment by replenishing the media every 3-4 days with fresh drug. For re-sensitization experiments, persister populations were derived by sorting day 14 osimertinib-treated PC9-watermelon cells based on mCherry expression following 11 days of doxycycline chase. Following sorting, persister populations were propagated in drug free media for 20 passages prior to starting the assay. Cell numbers following treatment were determined by imaging plates and quantifying nuclei using the Acumen Cellista plate cytometer (TTP Labtech).

**Live cell imaging**

Cells were grown in poly-D-lysine-coated glass bottom plates (MatTek Corporation) and imaged using a Nikon Eclipse TE-2000 inverted microscope with a 10X Plan Apo objective and a Hammamatsu Orca ER camera, equipped with environmental chamber controlling temperature, atmosphere (5% $CO_2$) and humidity. For long-term live cell imaging experiments, drug containing media was replaced every day to prevent changes in drug concentration due to evaporation. Images were acquired using the MetaMorph Software. For the initial 24h images were acquired every 10 min, for 24-120h images were acquired every 20 min and later images were acquired hourly.

Single-Cell Tracking was performed as in Reyes *et al.* 2018(Reyes et al., 2018). In brief, we used a semi-automated MATLAB based method to track and annotate cell fates. The method relies on identification of cell centroids using intensity and shape information of a constitutively expressed nuclear marker (H2B-mCherry), centroid linkage using nearest-neighbor, and user correction and annotation of cell fate events (Tracking software available at: https://github.com/balvahal/p53CinemaManual). A single cell of each progeny was tracked throughout the time course or until cell death. To determine the progeny size of each persister, cells were manually tracked cell death was manually recorded.

For doubling time assays, cells were seeded in 96-well plates at a density of $3\times10^3$ cells/well and measured using an IncuCyte ZOOM live cell imaging system (Essen BioScience, Ann Arbor, MI, USA).

**Cloning of the watermelon library**

A lentivirus backbone was constructed containing: TRE3GS promoter for H2B-mCherry and hPGK promoter for expression of Tet-On rtTA element, NLS-mNeon and a polyadenylated lineage barcode cassette. We prepared the vector backbone by digesting 20 µg of it with Sbf1 (New England Biolabs (NEB)) overnight at 37°C followed by gel purification using 1% E-Gel Ex (ThermoFisher Scientific). The cut backbone was extracted with the QIAprep miniprep Kit (Qiagen) and the resulting DNA was eluted in 100 µl of $H_2O$ and purified with 0.70× AMPure XP SPRI beads.

The double-stranded lineage barcodes were generated by annealing two DNA primers, a 90-bp-long oligonucleotide containing a semi-random 30-bp-long barcode sequence (*i.e.*, 15 repeats of A/T (W)–G/C (S)) and a flanking primer pair for barcode amplification and preforming a single cycle extension reaction (for primer list see **Supplementary Table 3**). The resulting oligonucleotides were ran on 2% E-Gel Ex and purified with 2.5×AMPure XP SPRI beads.

To generate a watermelon library with a diverse lineage barcode pool, the lineage oligonucleotides were mixed together with the backbone fragment in a 5:1 molar ratio together with an equal volume of Gibson master mix (Gibson Assembly Cloning Kit NEB), incubated at 50°C for 4 h, cleaned with 0.75× AMPure XP SPRI, eluted in 15 µL $H_2O$ and electroporated into Endura competent cells (Lucigen). We expanded the cells in LB liquid culture supplemented with carbenicillin (Sigma-Aldrich) for 16 hours at 30°C and purified the pooled library plasmid with the Endotoxin-Free Plasmid Maxiprep Kit (Qiagen). Library complexity was estimated by sequencing the library plasmid pool at a depth of approximately 68 million reads.

**Lentivirus production**

Lentiviral particles were produced by transfecting 293T cells with dVPR and VSVG packaging plasmids, using the X-tremeGene transfection reagent (Sigma-Aldrich) according to the manufacturer's instructions. Media was replaced with DMEM medium (ThermoFisher Scientific) supplemented with 20%FBS 20 hours post transfection, and media containing virus particles were collected 48 hours post transfection. For ORF overexpression and guide screen, virus particles were concentrated using Amicon 100 KDa 15mL columns (Millipore) in a cold centrifuge at 1,000 xg to a final concentration of 500 µl per virus. Virus was aliquoted and stored at −80°C until use.

**Watermelon cell line construction**

Parental cell lines were transduced using the watermelon virus and cells were spin infected using 16 µg/ml polybrene in 2,500 rpm for 30 min at 30°C. After a 24h incubation with virus, media was changed and 72 hours post infection cells were sorted for mNeon expression. To ensure that the majority of cells were labeled with a single barcode per cell, for watermelon lentiviral infection, we used a target multiplicity of infection (MOI) of at most 0.3, corresponding to less than 30%

mNeon expressing cells 72 hours post infection. Sorted cell populations,10,000 cells each, were expanded in culture for three passages, aliquoted to $2X10^6$ cells per vial and stored in liquid nitrogen.

**Single-cell capture for time course experiment**

Watermelon cells were thawed, expended in dox (2μg/ml) containing media for 96 hours and mCherry positive cells were sorted using a MoFlo Astrios Cell Sorter (Beckman Coulter) and re-plated. Following 96 hours of recovery, the cells were seeded into six-well plates at 300,000 cell per well and were given 24h to attach prior to adding 300nM osimertinib. Dox was continuously added to the media until day 3 of drug treatment. Cells were harvested at day 0 (untreated), 3, 7 and 14 of drug treatment. To obtain cell suspension for single cell profiling, cells were scrapped from the well, washed and resuspended in FACS buffer (0.5% BSA in phosphate-buffered saline), and filtered through a 40μm strainer. To delineate the differences between persister populations, day 14 cells were gated based on mCherry expression. Following sorting, the cells were spun down and approximately 9,000 single cells per sample were loaded to the Chromium Controller (10x Genomics). ScRNA-seq libraries were generated using the 10X Genomics Chromium Single Cell 3' Kit v2 and the 10x Chromium Controller (10x Genomics) according to the standard v2 protocol. The resulting 3' scRNA-Seq libraries were pooled together and sequenced with a HiSeq (Illumina, R2 read length 98 base pairs). To increase lineage barcode capture, targeted sequencing of the barcode area was performed using the whole transcriptome amplification product generated as a part the v2 protocol as a PCR template (for primer list see **Supplementary Table 3**). Targeted libraries were gel purified and sequenced with a MiSeq (Illumina).

**RNA-seq data analysis**

*Read alignment and data processing*

Reads were mapped to the GRCh38 human transcriptome using CellRanger 2.1.0 (10x Genomics), and transcript-per-million (TPM) was calculated for each gene in each filtered cell barcodes sample. TPM values were then divided by 10 (TP100K), since the complexity of our single-cell libraries is estimated to be on the order of 100,000 transcripts. For each cell, we quantified the number of genes expressed and the proportion of the transcript counts derived from mitochondrial encoded genes. Cells with either <1,000 or more than 4,200 detected genes or >0.1 mitochondrial fraction were excluded from further analysis. Finally, the resulting expression matrix was filtered to remove genes detected in <3 cells. All the above steps were done using the Seurat v2 R package(Stuart et al., 2019).

*Detection of differentially expressed gene signatures*

To identify cellular programs that are associated the ability of persisters to cycle, we searched for differentially expressed, cell cycle independent, gene signatures that show the largest difference between the persister subpopulation. First, we identified genes are differentially expressed (had an adjusted P value lower than 0.001 and a |log2FC|>0.2) in the cycling persisters after regressing out known cell-cycle genes based on a published gene list(Tirosh et al., 2016) using the MASTDETest implemented in Seurat. Next, we used hypergeometric to test which gene signatures were enriched in this gene set. This resulted in 37 gene signatures (FDR q-value<0.001). Last, for each of the 37 gene signatures, we calculated an Overall Expression signature score per cell, as previously described(Jerby-Arnon et al., 2018) and the mean Overall Expression signature score per sample. Finally, signatures in which the mean signature level was lower for persister cells compared with untreated day 0 cells, were filtered out (see **Supplementary Table 1** for final signature scores).

*Force directed layout embedding*

We first identified variable genes using Seurat's FindVariableGenes function (mean.function = ExpMean, dispersion.function = LogVMR). The log-transformed gene expression matrix of the 1,297 variable genes was used to compute a 20-dimensional diffusion component embedding using the function DiffusionMap (sigma = "local",k=1422,n_eigs=20) implemented in the Destiny R package(Angerer et al., 2016) .The resulting 20-dimensional diffusion component space was used to build a *k*-nearest neighbor graph (*k*=15) using the nng function from the cccd R package. To generate 2D visualization of the resulting graph, we applied a force-directed layout embedding on the graph using the ForceAtlas2 algorithm(Jacomy et al., 2014) from the Gephi Toolkit (v0.9.2).

*Mapping cell lineage*

To increase lineage detection rate, lineage barcode PCR dial-out, rather than 10x whole transcriptome sequences were used for generating a cell-barcode (CB) lineage-barcode (LB) map. First, dial out sequences were filtered based on CellRanger's "barcodes.tsv" file to contain only valid CB reads. Second, CB reads were removed if they didn't follow the expected semi-random sequence pattern (15 repetitions of A or T followed by a C or G). The retained sequences were used to generate a CB-LB table. We next calculated how many UMIs support each CB-LB pair. For each cell barcode, we calculated the Levenshtein edit distances of all its associated lineage barcodes and collapsed all LB that were within an edit distance of 3 from the most common LB of this cell. If after collapsing, the less prevalent LB were supported by only one UMI, they were removed. As an additional filtering step, LB that were associated with the cell less than three times compared to the most supported LB were also removed as they were likely introduced by the PCR amplification. Finally, we retained only CB-LB that were supported by at least two UMIs.

*Mapping lineage fate*

Lineages were assigned one of four fates: "drug-sensitive/absent on day 14", "cycling", "non-cycling", or "multi-fate". "Drug-sensitive/absent on day 14" lineages were defined as those lineages that were not detected at day 14 (lack of detection is likely due to drug sensitivity but be cannot rule out that some clones are absent due to sampling). Cycling lineages were defined as those lineages in which all progeny belonged to mCherry$^{low}$ or mCherry$^{medium}$ sample in day 14. Non-cycling lineages were lineages in which all cells belonged to the mCherry$^{high}$ sample in day 14. Multi-fate lineages were lineages in which cells were found both in the mCherry$^{low}$ or mCherry$^{medium}$ samples and in the mCherry$^{high}$ sample in day 14. To test if multi-fate lineages were less frequent than expected by chance, we performed a permutation test. We shuffled the assignment of cells to lineage barcode of day 14 mCherry assignment group, keeping the overall number of cells in each group as in the original dataset. We next assigned each lineage its fate (cycling, non-cycling, or multi-fate) based on the randomized cell identity. This process was repeated 1,000 times to generate a distribution based on the number of multi-fate lineages observed in each iteration.

*Correlation between persister lineage size and gene expression*

To identify genes that are associated with increased lineage size, we correlated the expression of each of the variable genes with day 14 clone size. We first filtered the expression matrix to contain only cells in which a lineage barcode was identified. We next filtered genes that in a given time point had no variance. For each time point, we calculated the Pearson correlation between the expression of each gene across all cells with the vector of day 14 lineage size. This analysis resulted in a single correlation value for each gene.

*Calculating Overall Expression signature scores*

Given a gene signature and a gene expression matrix, we first binned the genes to 10-50 expression bins (depending on data complexity) according to their average expression across the cells or samples. For each gene signature, we sampled 100 random compatible signatures for normalization. The final reported score is computed by subtracting the 'real' mean signature score from the randomized one. For more detailed description of the method please refer to Jerby-Arnon, L. *et al.(Jerby-Arnon et al., 2018)*

**Metabolite profiling**

Polar cell extracts were profiled using negative and positive ionization mode using liquid chromatography tandem mass spectrometry (LC-MS) methods. Negative ionization mode data were acquired using an ACQUITY UPLC (Waters Corp, Milford MA) coupled to a 5500 QTRAP triple quadrupole mass spectrometer (AB SCIEX, Framingham MA). Positive ionization data were acquired using an LC-MS system composed of a Shimadzu Nexera X2 U-HPLC (Shimadzu Corp) coupled to a Q Exactive hybrid quadrupole orbitrap mass spectrometer (ThermoFisher Scientific). For both modes, cell-sorted samples were extracted using 200 µl 80% methanol containing 0.5 ng/µL inosine-$^{15}N_4$, 0.5 ng/µL thymine-$d_4$, and 1 ng/µL glycocholate-$d_4$ as internal standards (Cambridge Isotope Laboratories, Inc., Tewksbury MA).

For the negative extraction, 90µl of each sample was centrifuged (10 min, 9,000 x *g*, 4°C) and the supernatants (10 µL) were injected directly onto a 150 x 2.0 mm Luna NH2 column (Phenomenex, Torrance CA). The column was eluted at a flow rate of 400 µL/min with initial conditions of 10%

mobile phase A (20 mM ammonium acetate and 20 mM ammonium hydroxide (Sigma-Aldrich) in water (VWR)) and 90% mobile phase B (10 mM ammonium hydroxide in 75:25 v/v acetonitrile/methanol (VWR)) followed by a 10 min linear gradient to 100% mobile phase A. The ion spray voltage was -4.5 kV and the source temperature was 500°C. Raw data were processed using MultiQuant 3.0.3 software (AB SCIEX, Framingham MA) for automated peak integration.

For the positive extraction, 80µL of each sample was dried down using the turbovap (TurboVap LV, Caliper Life Sciences), Each sample was resuspended in 8µL of water and then crash with 72µL extraction solution 74.9:24.9:0.2 v/v/v acetonitrile/methanol/formic acid containing stable isotope-labeled internal standards (valine-d8, Sigma-Aldrich; and phenylalanine-d8, Cambridge Isotope Laboratories). Extracts were vortexed for 1 minute, samples were spun at 10,000 rcf for 10 minutes at 4°C and the resulting supernatant was moved to autosampler vials. Raw data were processed using TraceFinder software (Thermo Fisher Scientific) and Progenesis QI (Nonlinear Dynamics).

**Metabolomics data analysis**

Relative abundance metabolite data from positive and negative ionization mode extractions were analyzed in-part using the MetaboAnalystR package(Chong et al., 2019). First, metabolites containing any missing values across samples were removed from the dataset. For metabolites that were detected in both positive and negative ionization extractions, we only considered values from the negative extraction as it used a triple quad for the analysis, which often yields higher accuracy. Prior to statistical analysis, the values of each metabolite across samples were log-normalized and mean-centered. Significantly different metabolites detected by ANOVA were clustered by complete linkage hierarchical clustering and visualized in heatmap format using the pheatmap R package.

**Fatty acid oxidation measurements**

For FAO assays, cells in 6 well-plates were treated with 100µM etomoxir as indicated. Pulsing was performed in serum-free medium containing 1mM carnitine with 0.75 µCi [9,10(n)-3H] palmitic acid (GE Healthcare) for 2 hours. The medium was collected and eluted in columns packed with DOWEX 1X2-400 ion exchange resin (Sigma) to analyze the released $^{3}$H2O. $^{3}$H2O was measured in counts per minute (CPM) and normalized to total cellular protein using BCA Protein Assay Kit (ThermoFisher Scientific).

**Reactive oxygen species measurements**

To measure relative levels of reactive oxygen species, drug treated and control cells were stained with 5 μM CellROX Deep Red Reagent (Thermo Fisher) for 30 min at 37°C, washed three times with PBS, trypsinized and analyzed using MoFlo Astrios Cell Sorter (Beckman Coulter).

**ORF assays**

Purified Open Reading Frame (ORF) expression vectors for CPT1A and GPX2 were ordered from Genecopoeia (CPT1A: EX-A1436-Lv156, GPX2: EX-A3079-Lv156). PC9 cells harboring the watermelon construct were transduced with individual vectors and treated with drug as previously described. To quantify the increase in expression. We confirmed increased expression of each ORF by reverse transcription and quantitative polymerase chain reaction (RT-qPCR). RT-qPCR primers were designed to: (1) capture both endogenous and plasmid sequences of each gene; (2) span only gene introns; (3) produce amplicons no larger than 200bp; and (4) have melting temperatures of 60±1˚C (for primer list see **Supplementary Table 3**). For each sample, two replicates of 20,000 cells each were lysed in RLT buffer (Qiagen #79216) and RNA was isolated using Dynabeads MyOne Silane beads (ThermoFisher Scientific #37002D). Next, samples were treated with TURBO DNase (ThermoFisher Scientific #AM2238), and subsequently purified again with silane beads. Reverse transcription was preformed using SuperScript II Reverse Transcriptase (ThermoFisher Scientific #18064022) according to manufacturer's protocol. The resultant cDNA was diluted 1:10 and qPCR was performed with SYBR Green Master Mix (ThermoFisher Scientific #4368706).

**Patient data analysis**

**Lung**

Lung cancer scRNA data was obtained from https://github.com/czbiohub/scell_lung_Adenocarcinoma. To ensure that only epithelial cancer cells would be included in our analysis, we used the script "NI04_Cancer_cells_DEgenes.Rmd" provided in the github repository to filter for tumor cells (using: “inferCNV_annotation==tumor”) this resulted in 3,620 cells. As our study main focus is EGFR-driven lung cancer, we further filtered the data to include only samples where EGFR was considered to be the main cancer driver mutation (using: subset=driver_gene=="EGFR"). This resulted in 2,020 EGFR-driven cancer cells of which 426, 500, and 1,096 belonged to treatment naïve (TN), residual disease (RD) and progressive disease (PD) groups, respectively (grouping was based on the “analysis” column in the metadata, see **Supplementary Table 2** for additional sample information). Next, the data were normalized and scaled and the cells were scored for cell cycle, fatty acid metabolism (FAM) and reactive oxygen species (ROS) signatures (a full list of signatures used in this study is in **Supplementary Table 4**). Wilcoxon test was used to estimate if the treatment groups were significantly different from each other. We next assigned the cells as either cycling or non-cycling using Seurat’s built-in “CellCycleScoring” function. Cells were classified as cycling if they were assigned as “G2M” or “S” phase by the algorithm, and non-cycling if they were classified as being in the “G1” phase. To account for patient and cell complexity effects (as cycling cells are expected to have higher overall number of transcripts), we performed a mixed effect logistic regression using the lmer function of the lmerTest package in R, with the model signature score ~ nCount_RNA + cell_fate + (1|sample_name), allowing the intercept to vary by sample. P values were calculated using Satterthwaite's method for denominator degrees of freedom.

##### Melanoma

Melanoma bulk RNA-seq data was downloaded from https://www.ebi.ac.uk/ega/dacs/EGAC00001000324. To quantify how treatment affected signature induction, we first filtered the dataset to contain only the 11 patients where both pre-treatment and run-in samples were available. For these samples (**Supplementary Table 2**), we calculated the log 2-fold change in reads per kilobase of transcript, per million mapped reads (RPKM) for each of the genes in the genome. This was done by dividing the RPKM measured in on-treatment sample by the RPKM measured in the pre-treatment sample. Finally, we calculated the median signature score for cell cycle, FAM and ROS. To obtain pre-treatment signature scores, we calculated a median signature score using the RPKM values only from the pre-treated samples of the 11 patients without any division.

**Breast**

Breast cancer microarray data was downloaded from https://www.ncbi.nlm.nih.gov/geo/query/acc.cgi?acc=GSE130786 and https://www.ncbi.nlm.nih.gov/geo/query/acc.cgi?acc=GSE130787. To quantify how lapatinib treatment affects signature induction, we first filtered the dataset to contain only the 18 patients from the lapatinib arm ("TCTy" group) where both pre-treatment and run-in samples were available (**Supplementary Table 2**). To remove noise from the data, we used the genefilter package in R to filtered out probes that had an average expression within the $25^{th}$ expression percentile in at least 80% of the samples. For each gene, we than took the probe with the highest absolute intensity value. As microarray values already represent the ratio between two conditions, to calculate changes in signatures following treatment, we transformed the values to be on a log-2 scale and calculated the median of FAM and ROS signature for each patient. Baseline signature values were calculated using the set of microarrays where each sample was compared to a breast tumor mixed reference pool.

**Data availability**

RNA-seq data have been deposited in the NCBI Genome Expression Omnibus (GEO) under the accession code GSE150949. Watermelon plasmid will be deposited to Addgene upon acceptance of the manuscript.

**Statistical analyses**

Statistical tests and graphing of data were performed with GraphPad Prism v.7.0a and R v. 3.6.1. Unless otherwise noted, P values were calculated using unpaired, two-tailed t-tests assuming unequal variance. Multiple hypothesis correction was done using Holm's method.

#### Supplementary Figures

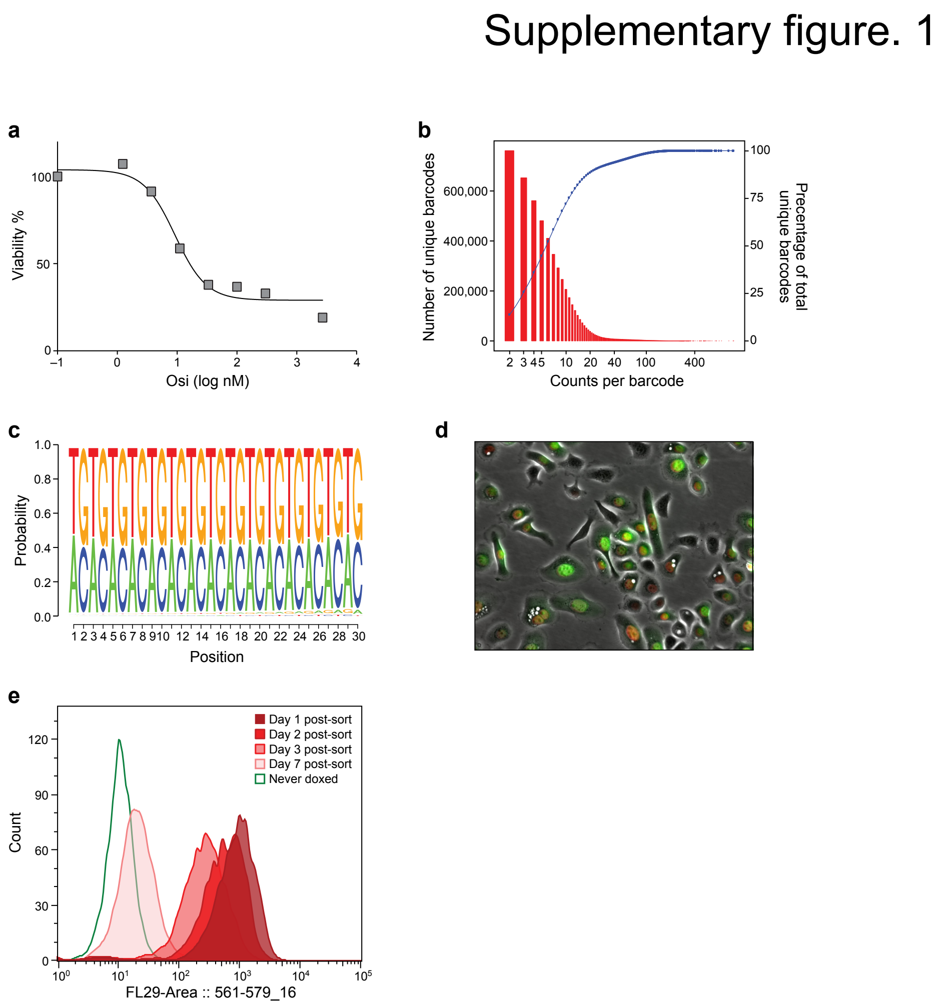


**Supplementary Figure 1. Lineage detection efficacy and fluorescent dilution capacity of the Watermelon library**

***a.*** Viability of PC9 cells treated osimertinib. % of viable PC9 cells (*y* axis) after 72hr of treatment with osimertinib at different concentrations (*x* axis). ***b.*** Watermelon library complexity. Distribution of number of unique lineage barcodes (*y* axis, red bars) in a Watermelon plasmid library sequenced at a depth of ~68X$10^6$ reads. Blue curve: cumulative wealth distribution of unique barcodes. **c.** Watermelon library sequence diversity. Sequence logo of nucleotide composition at each position (*x* axis) relative to the beginning of the barcode sequence of 5,472,944 unique lineage barcodes detected in the Watermelon library. ***d.*** PC9-Watermelon cell line grown in dox containing media. A merge of the green, red and bright field channels is shown. Scale bar

20μm. ***e.*** Fluorescence dilution of H2B-mCherry over time reports proliferative history. Distributions of mCherry fluorescence level (*x* axis) for *n*=3000 cells analyzed by flow cytometry at each time point (color legend) from cells transduced with the Watermelon vector, exposed to dox for 48 hours, sorted for red positive cells and seeded in separate wells at t=0 (**Methods**). Data are representative of two independent experiments.

### Supplementary figure. 2

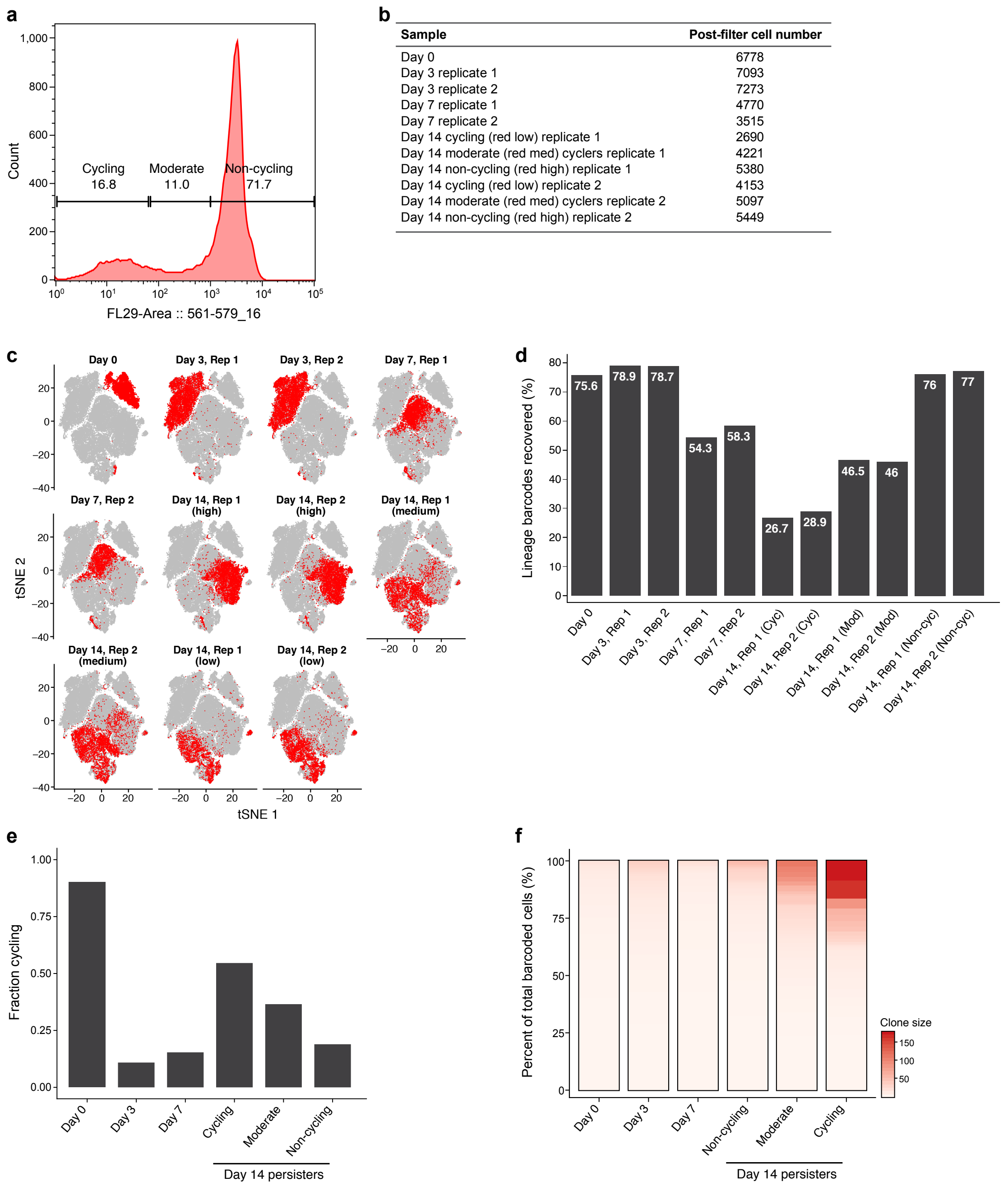


**Supplementary Figure 2. scRNA-Seq along a time course of osimertinib treated PC9-Watermelon cells**

***a.*** Sorting strategy. Distribution of mCherry fluorescence level (x axis) in Watermelon-PC9 cells gates at day 14 of osimertinib treatment, marked by representative sorting gates using to sort three persister subpopulations: cycling, moderate cyclers and non-cycling. ***b.*** Number of high-quality cells profiled in each sample. ***c.*** Changes in expression profiles following treatment. *t*-stochastic neighborhood embedding (tSNE) of 56,419 PC9-Watermelon cell profiles (dots), colored (red) by the labeled time point. d. Assignment of cells to lineages by lineage barcode. Percent of cells (y axis) at each time point/subpopulation (*x* axis) that have a detected lineage barcode. ***e.*** Identification of cycling cells. Percent of cells (*y* axis) at each time point/subpopulation (*x* axis) that express either the G2/M or S phase signature. ***f.*** Single cell-derived clone size by sample. In each sample, detected barcodes were sorted in descending order by the sum of their counts. Each unique lineage barcode was accounted as a separate clone.

### Supplementary figure. 3

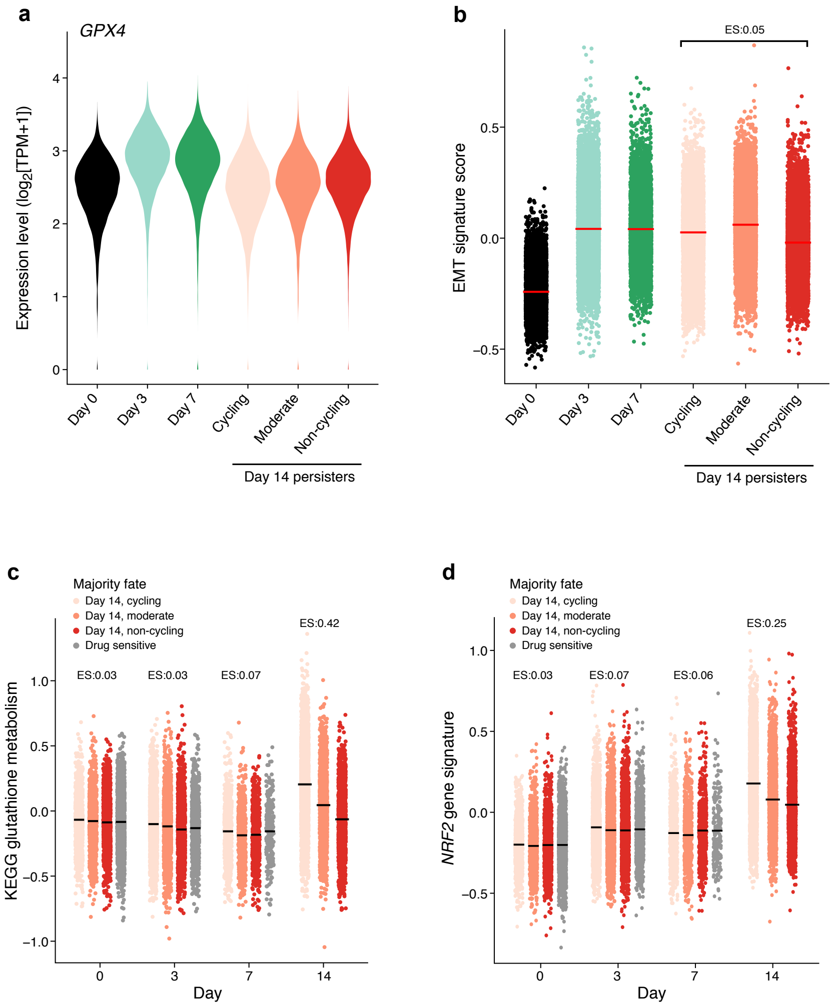


**Supplementary Figure 3**. **Differences in transcriptional programs in cycling and non-cycling persisters**

***a,b.*** GPX4 and EMT signature expression are similar in cycling and non-cycling persisters. Distribution of expression levels of GPX4 (*y* axis, $\log_2$(TPM+1), a) and EMT signature (*y* axis, b) across time points and subpopulations. Effect size (ES, b): difference between the mean signature score of cycling and non-cycling persisters. ***c,d.*** Higher expression of glutathione metabolism and NRF2 signatures in cycling *vs*. non-cycling persisters. Signature score (*y* axis) of glutathione metabolism (c) and NRF2 pathway (d) signatures in cells profiled at each time point (*x* axis) stratified by their lineage majority fate at day 14 (color legend). Effect size (ES) indicates difference between the mean signature score of cycling and non-cycling persisters.

Supplementary figure. 4

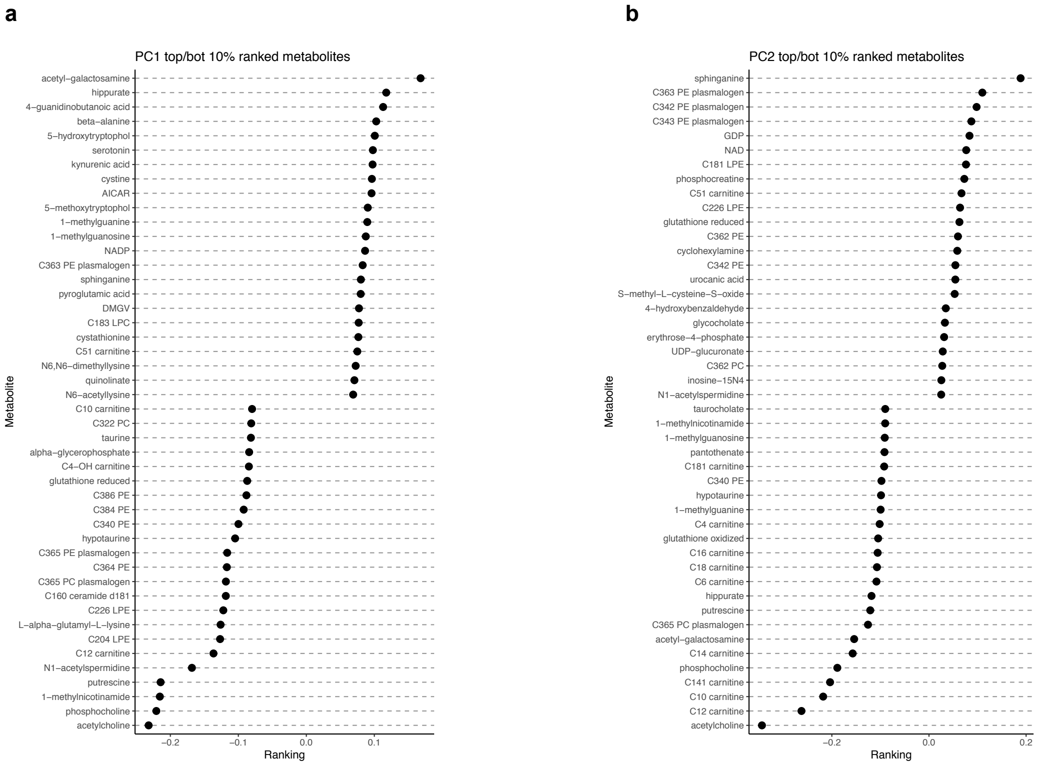


**Supplementary Figure 4. Principal Component Analysis (PCA) of metabolite profiles of cycling persisters, non-cycling persisters and untreated parental cells.**

Loadings (*x* axis) for the top 46 metabolites (y axis) associated with PC1 (a) and PC2 (b).

Supplementary figure. 5

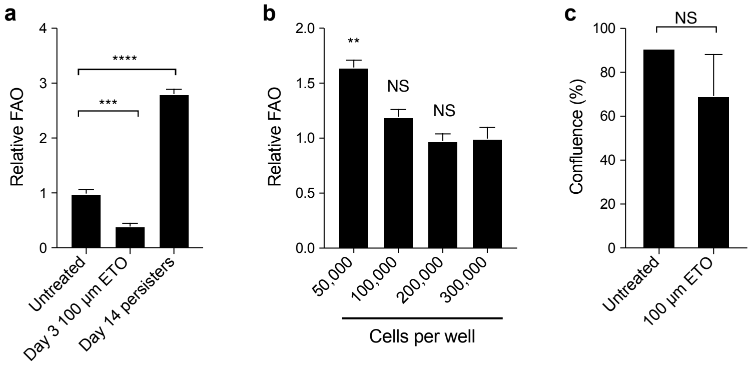


**Supplementary Figure 5. Effect of drug treatment and cell density on FAO measurement**
***a.*** Mean fatty acid oxidation (FAO) level (y axis, relative to mean of the untreated controls) measured by 3H-palmitic acid oxidation in PC9-Watermelon cells either untreated, treated only with 100 µM etomoxir for 3 days, or treated with 300nM osimertinib for 14 days. ***b.*** Mean FAO levels (*y* axis, relative to cells seeded at 300,000 per well, as used for the osimertinib time course) in PC9-Watermelon cells seeded at different densities (*x* axis) 24 hours prior to measurement. ** P< 0.01, two tailed *t*-tests; NS – not significant (compared to 300,000 cells per well). ***c.*** Mean confluence (*y* axis) of PC9-Watermelon cells treated with 100µM Etomoxir for 14 days (**Methods**). Error bars: SD. n=2-3.

### Supplementary figure. 6

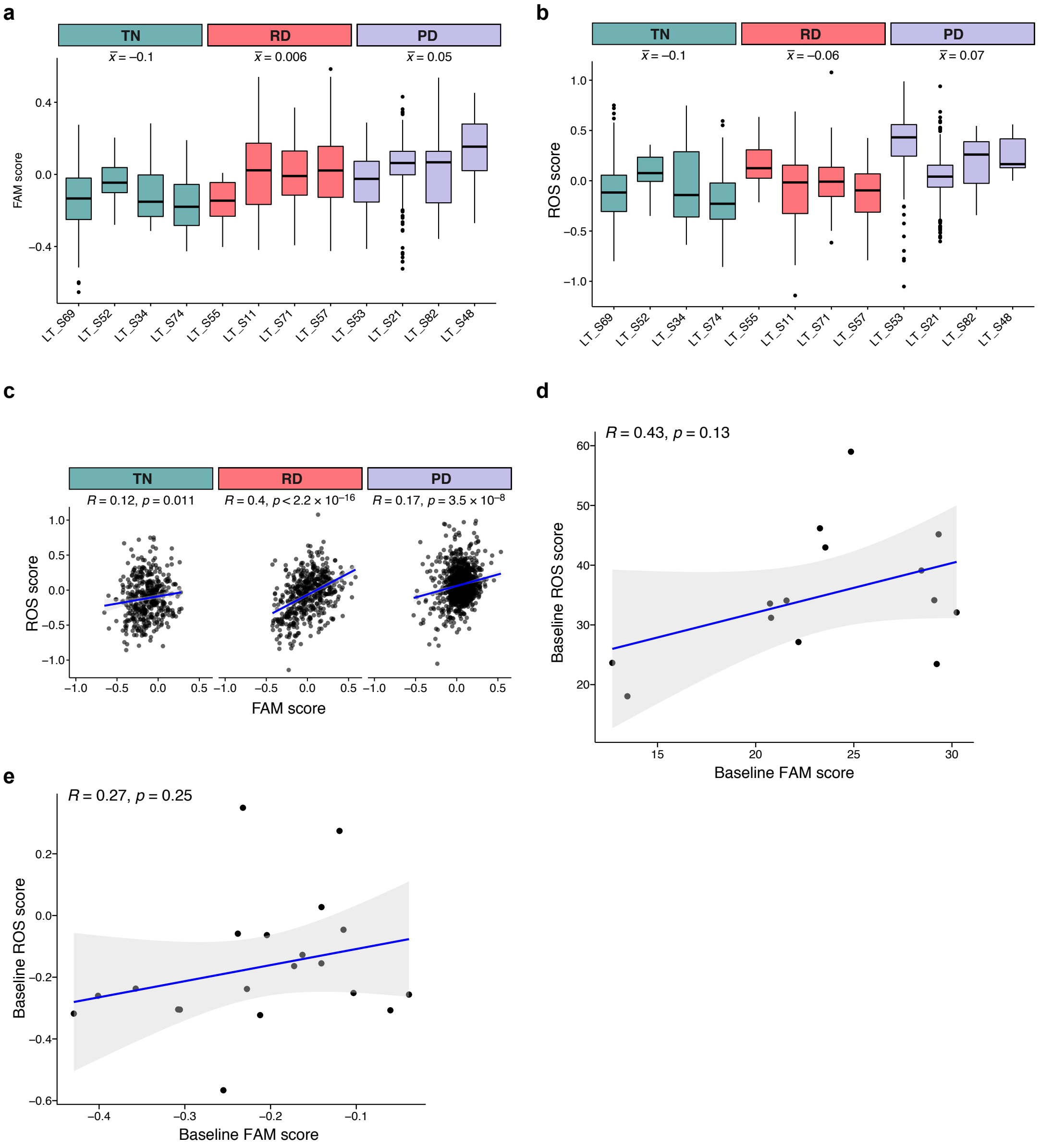


**Supplementary Figure 6. Changes in expression of metabolic programs in patient tumors**
**a,b.** Increase in fatty acid metabolism and ROS pathway signatures in drug-treated human lung adenocarcinoma. Distribution of expression scores of FAM (a) and ROS (b) signatures in cells from individual EGFR-driven lung adenocarcinoma tumors (with more than 10 cells) across different treatment timepoints (*x* axis). x̄: Mean signature level for time point. ***c-e.*** Correlation between ROS (y axis) and FAM (x axis) signature scores in (c) treatment naïve (TN), residual

disease (RD) and progressive disease (PD) human lung adenocarcinoma, (d) treatment naïve melanoma, and (e) treatment naïve breast cancer.

**Supplementary Tables**

**Supplementary Table 1.** Mean signatures expression of the three different persister subpopulations

**Supplementary Table 2.** Patient information

**Supplementary Table 3.** Primers used in this study

**Supplementary Table 4.** Gene signature list
